# Learning to embed lifetime social behavior from interaction dynamics

**DOI:** 10.1101/2021.09.01.458538

**Authors:** Benjamin Wild, David M Dormagen, Michael L Smith, Tim Landgraf

## Abstract

Interactions of individuals in complex social systems give rise to emergent behaviors at the group level. Identifying the functional role that individuals take in the group at a specific time facilitates understanding the dynamics of these emergent processes. An individual’s behavior at a given time can be partially inferred by common factors, such as age, but internal and external factors also substantially influence behavior, making it difficult to disentangle common development from individuality. Here we show that such dependencies on common factors can be used as an implicit bias to learn a temporally consistent representation of a functional role from social interaction networks. Using a unique dataset containing lifetime trajectories of multiple generations of individually-marked honey bees in two colonies, we propose a new temporal matrix factorization model that jointly learns the average developmental path and structured variations of individuals in the social network over their entire lives. Our method yields inherently interpretable embeddings that are biologically relevant and consistent over time, allowing one to compare individuals’ functional roles regardless of when or in which colony they lived. Our method provides a quantitative framework for understanding behavioral heterogeneity in complex social systems, and is applicable to fields such as behavioral biology, social sciences, neuroscience, and information science.

**Author summary:** Group-level emergent behaviors are the result of interactions between individual group members. To understand these social dynamics, one must objectively measure the function of an individual in their group at any given time. Ideally, one would also like to compare individuals from different groups, for example, to measure how specific environmental conditions or other external factors influence group behavior. Unfortunately, such an objective measure is hard to obtain because the group and its dynamics constantly change, making it challenging to define an individual’s role in the group as a function of its actions and interactions. We propose a principled approach to model individuals in complex social systems by considering that function often depends, at least partially, on common factors such as age. The model learns a meaningful and interpretable descriptor for all individuals, and can be used to understand how complex social systems function and the emergence of group behavior.

## 1 Introduction

Animals living in groups often coordinate their behavior, resulting in emergent properties at the group level. The dynamics of the inter-individual interactions produce, for example, the coherent motion patterns of flocking birds and shoaling fish, or the results of democratic elections in human societies. In many social systems, individuals differ consistently in how, when, and with whom they interact. The way an individual participates in social interactions and therefore contributes to the emergence of group-level properties can be understood as its functional role within the collective [1–4].

Technological advances have made it possible to track all individuals and their interactions, ranging from social insects to primate groups [5–10]. These methods produce datasets that have unprecedented scale and complexity, but identifying and understanding the functional roles of the individuals within their groups has emerged as a new and challenging problem in itself. Social network analysis of interaction networks has proven to be a promising approach because interaction networks are comparatively straightforward to obtain from tracking data, and the networks represent each individual in the global context of the group [2, 3, 11, 12].

In most social systems, the way individuals interact changes over time, due to new experiences, environmental changes, or physiological conditions. Furthermore, groups themselves also tend to change, both in size and composition [13–18]. Despite these changes over time, an objective measure of the functional role should identify individuals that serve a similar function (e.g. a guard versus a forager). Unfortunately, we are now facing a recursive definition of function: We are trying to derive the function of an individual from the network, but the network itself is also a function of the individuals’ behavior (and other factors). Still, consider a group-living species in which only a subset of individuals engage in nursing duties. If we analyze the networks of different groups of the same species in different environmental conditions and group sizes, we still expect an objective measure of function to be shared among individuals engaged in nursing, regardless of these confounding factors. How can we extract such an objective measure from a constantly changing network of interactions without a fixed frame of reference?

In many social systems, individuals share common factors that partially determine the roles they take. For example, an individual’s age can have a strong influence on behavior. In humans, factors such as socioeconomic status are comparatively easy to measure yet determine behavior and, therefore, interactions to a large extent. If individuals take on roles partially determined by a common factor, can we use this dependency to learn an objective measure of function? Here, we show that such common factors are a powerful inductive bias to learn semantically consistent functional descriptors of individuals over time, even in highly dynamic social systems.

In recent years, methods that automatically learn semantic embeddings from high-dimensional data have become popular. These methods map entities into a learned vector space. For example, in natural language models, a word can be represented as a vector, such that specific regions in the manifold of learned embeddings correspond to words with similar meaning. Similarly, recommender systems can learn meaningful embeddings of users and items, for example, movies, such that similar entities cluster in the manifold of learned embeddings [19–22].

Such embeddings are usually learned from the data without additional supervision. In recommender systems, a movie’s genre is usually not given in a dataset of user ratings, yet the genre can be identified given the learned embeddings [23]. This capability of learning embeddings from raw data and using them in downstream tasks is desirable in datasets of social interactions, where raw data is often abundant but labels are hard to acquire. Furthermore, embeddings are interpretable. For example, vector arithmetic of word embeddings can be used to understand how semantic concepts the natural language model has learned from the data relate to each other [24]. For entities that change over time, trajectories of embeddings can be analyzed, i.e., how one entity changes within the learned manifold of embeddings. Such analyses can, for example, reveal how environmental conditions such as resource availability affect behavioral changes within the group [25, 26].

Most real-world networks have a hierarchical organization with overlapping communities, and thus soft community detection algorithms are often used to group and describe entities [26–28]. Non-negative matrix factorization (NMF) is a principled and scalable method to learn embeddings from data that can be represented in matrix form, such as interaction networks. NMF has an inherent soft clustering property and is therefore well suited to derive embeddings from social interaction networks [29]. If the embeddings allow us to predict relevant behavioral properties, they serve our understanding as *semantic* representations.

In symmetric non-negative matrix factorization (SymNMF), the dot products of any two individuals’ embeddings (*factor vectors*) reconstruct their interaction affinity [30, 31], see Figure 1 a and b). However, this algorithm has no straightforward extension in temporal settings where the interaction matrices change over time. The interaction matrices at different time points can be factorized individually, but there is no guarantee that the embeddings stay semantically consistent over time. The dot product is permutation invariant, therefore factorization can result in different embeddings depending on the optimization method being used, or noise in the data. Consider the hypothetical case of two groups of animals of the same species with two tasks, guards and nurses. Factorizing the interaction matrices of both groups will likely reveal two clusters, but there is no guarantee that the same cluster will be assigned to the same task for both groups. The same problem can occur in the case of only one group with new animals emerging and some dying over time without any changes in the distribution of tasks on the group level. In this case, the embeddings are not semantically consistent over time. The prediction of relevant behavioral properties will deteriorate, and individuals cannot be meaningfully compared against each other.

**Figure 1.**
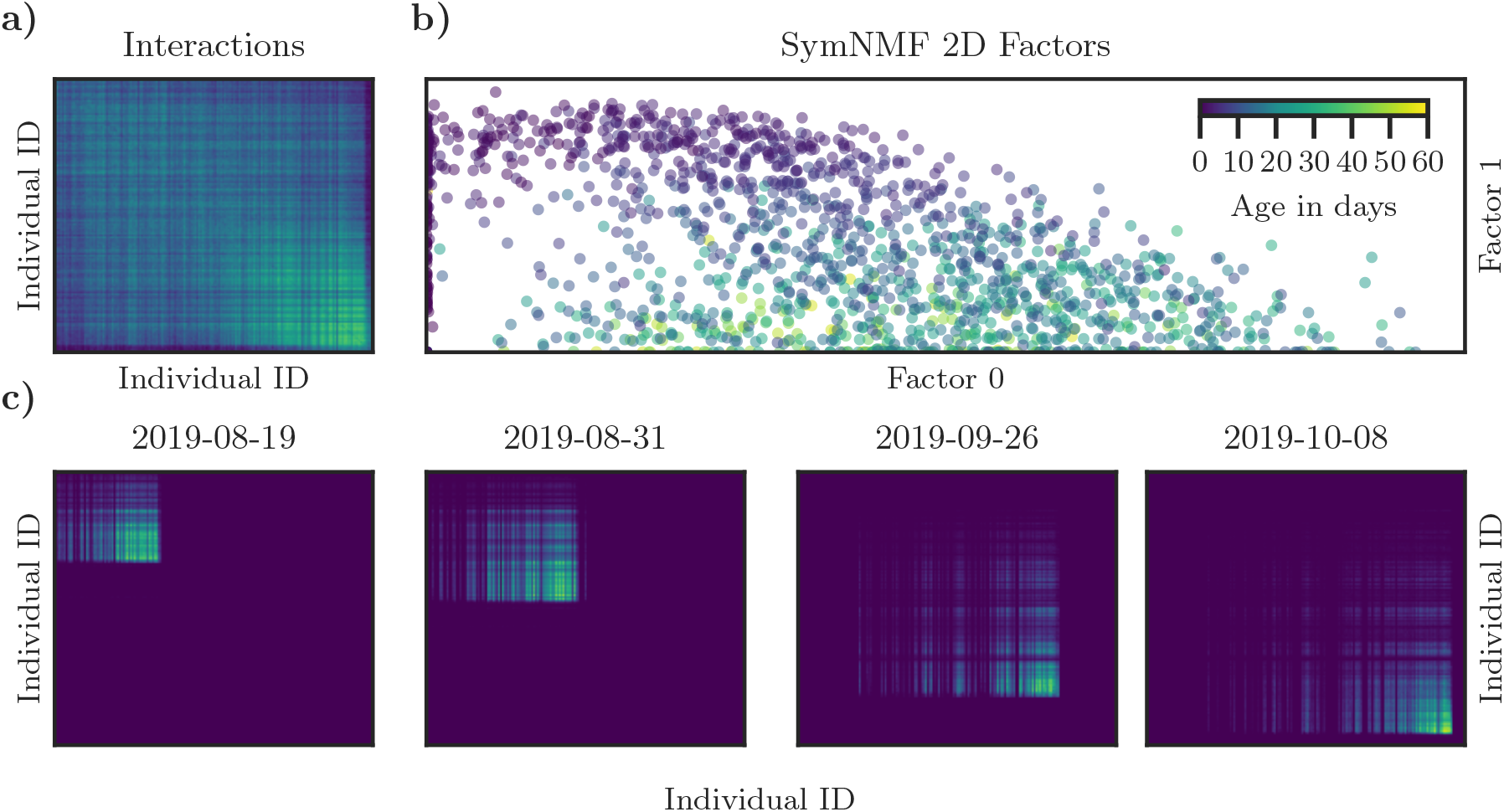
For a daily snapshot of a temporal social network, symmetric NMF is able to extract meaningful factor representations of the individuals. Colors represent the interaction frequencies of all individuals (**a**). The age-based division of labor in a honey bee colony is clearly reflected in the two factors - same-aged individuals are likely to interact with each other (**b**). For long observation windows spanning several weeks, the social network changes drastically as individuals are born, die, and switch tasks (**c**). Here, we investigate how a representation of temporal networks can be extracted, such that the factors representing individuals can be meaningfully compared over time, and even across datasets.

Several approaches to extend NMF to temporal settings have been proposed in a variety of problem settings. Previous work proposed factorization methods for time series analysis [32, 33], while others focus on the analysis of communities that are determined by their temporal activity patterns [34]. Jiao and coworkers consider the case of communities from graphs over time and enforce temporal consistency with an additional loss term [35]. Several previous works represent network embeddings as a function of time [36] and [37], but the meaning of these embeddings can still shift over time. Temporal matrix factorization is similar to the tensor decomposition problem, which has many proposed solutions, see review by [38]. In particular, time-shifted tensor decomposition methods have been used in multi-neuronal spike train analysis, when recordings of multiple trials from a population of neurons are available [39, 40].

We approach this problem in the honey bee, a popular model system for studying individual and collective behavior [41]. Honey bees allocate tasks across thousands of individuals without central control, using an age-based system: young bees care for brood, middle-aged bees perform within-nest labor, and old bees forage outside [42, 43]. While age is a good predictor for the task of an average bee, individuals often deviate drastically from this common developmental trajectory due to internal and external factors. Honey bee colonies are also organized spatially: brood is reared in the center, honey and pollen are stored at the periphery, and foragers offload nectar near the exit. Therefore, an individual’s role is partially reflected in its location, which provides the unique opportunity to evaluate whether learned embeddings based on the interaction data alone are meaningful.

A recent work proposes a method based on spectral decomposition to extract a semantic embedding (*Network age*) from honey bee interaction matrices and shows that these embeddings can be used to predict task allocation, survival, activity patterns, and future behavior [12]. The method proposed here is conceptually similar but solves several remaining challenges. Here, we introduce Temporal NMF (TNMF), which yields consistent semantic embeddings even for individuals from disjoint datasets, for example, data from different colonies, or for long-duration recordings that contain multiple lifetime generations.

TNMF jointly learns a) a functional form of the average trajectory of embeddings along the common factor, b) a set of possible functional deviations from the average trajectory, and c) for each individual, a soft-clustering assignment (*individuality embedding*) to these deviations. We show that these representations can be learned in an unsupervised fashion, using only interaction matrices of the individuals over time. We analyze how well the model is able to disentangle common development from individuality using a synthetic dataset. Furthermore, we introduce a unique dataset containing lifetime trajectories of multiple generations of individually-marked honey bees in two colonies. We evaluate how well the embeddings learned by TNMF capture the semantic differences of individual honey bee development by evaluating their predictiveness for different tasks and behaviorally relevant metrics compared to several baseline models proposed in previous works.

**Figure 2.**
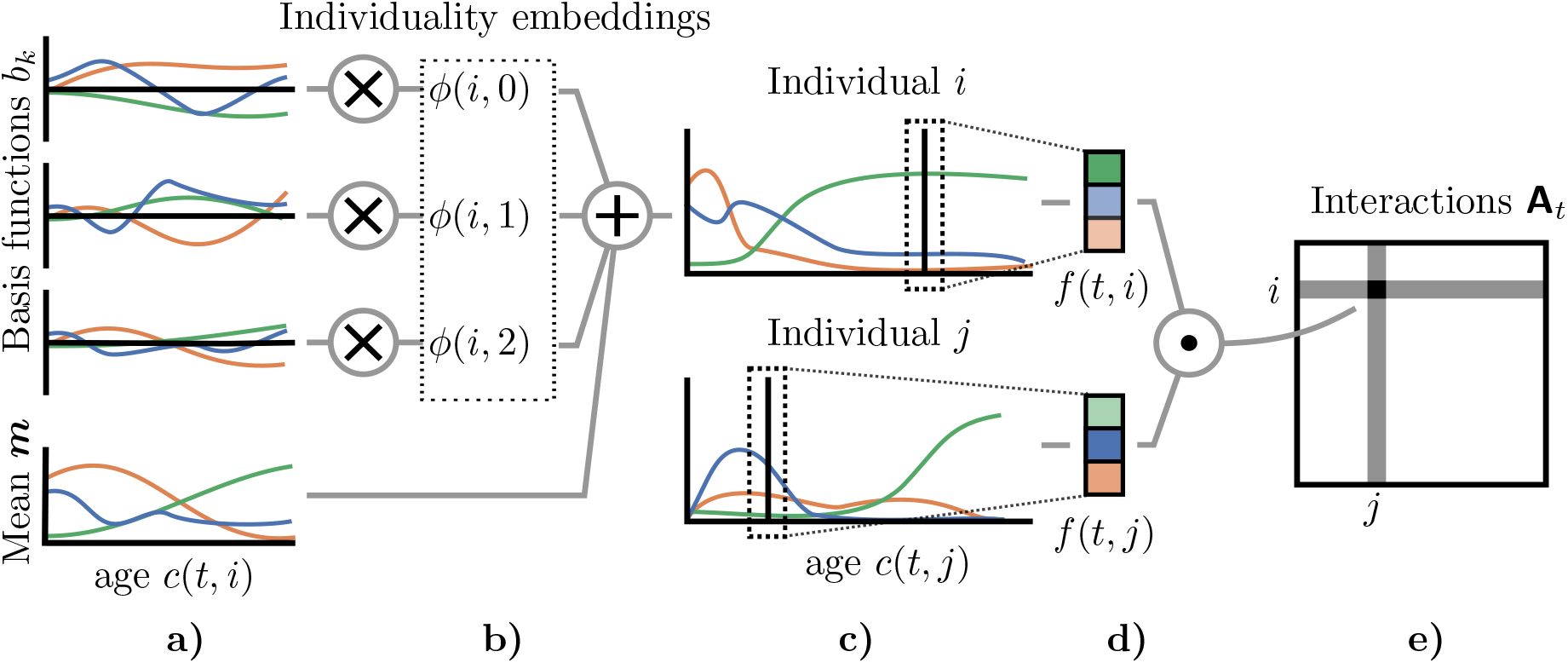
Overview of the method: We learn a parametric function describing the *mean life trajectory* ***m***(*c*(*t, i*)) and a set of basis functions of individual variation ***b***(*c*(*t, i*)), where *c*(*t, i*) is the age of individual *i* at time *t* (**a**). For each individual, an embedding is learned consisting of one scalar per basis function that scales the contribution of the respective basis function - this vector of weights makes up the *individuality embedding* of an individual (**b**). The mean trajectory ***m***(*c*(*t, i*)) plus a weighted sum of the basis functions ***b***(*c*(*t, i*)) constitute the *lifetime trajectory* of each individual (**c**). At each time point, factors can be extracted from the individual lifetime trajectories (**d**) to reconstruct the interaction affinity between individuals (**e**). Note that the lifetime trajectories are functions of the individuals’ ages, while interactions can occur at any time *t*.

## 2 Materials and Methods

### 2.1 Temporal NMF algorithm

SymNMF factorizes a matrix 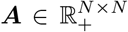 such that it can be approximated by the product ***FF***^*T*^, where 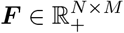 and *M* ≪ *N* :

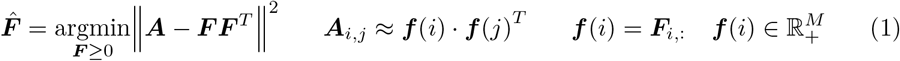

When applied to social networks, ***f***(*i*) can represent the role of an entity within the social network ***A*** [30, 31] - however, in temporal settings, factorizing the matrices for different times separately will result in semantically inconsistent factors.

Here we present a novel temporal NMF algorithm (*TNMF*) which extends SymNMF to temporal settings in which 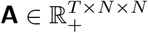 changes over time *t*. We assume that the entities *i* ∈ {0, 1, …, *N*} follow to some extent a common trajectory depending on an observable property (for example the age of an individual). We represent an entity at a specific point in time *t* using a factor vector ***f***^+^(*t, i*) such that

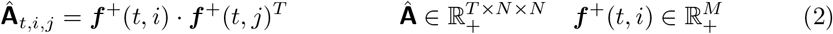

In contrast to SymNMF, we do not directly factorize **A**_*t*_ to find the optimal factors that reconstruct the matrices. Instead, we decompose the problem into learning an average trajectory of factors ***m***(*c*(*t, i*)) and structured variations from this trajectory ***o***(*t, i*) that depend on the observable property *c*(*t, i*):

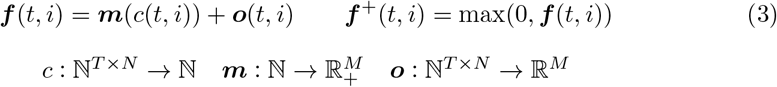

This decomposition is an inductive bias that allows the model to learn semantically consistent factors for entities, even if they do not share any data points (e.g., there is no overlap in their interaction partners), as long as the relationship between functional role and *c*(*t, i*) is stable. Note that in the simplest case *c*(*t, i*) = *t, TNMF* can be seen as a tensor decomposition model, i.e. the trajectory of all entities is aligned with the temporal dimension *t* of **A**. In our case, *c*(*t, i*) maps to the age of individual *i* at time *t*.

While many parameterizations for the function ***o***(*t, i*) are possible, we only consider one particular case in this work: We learn a set of *individuality basis functions* ***b***(*c*(*t, i*)) (shared among all entities) that define a coordinate system of possible individual variations and the *individuality embeddings ϕ*, which capture to what extent each basis function applies to an entity:

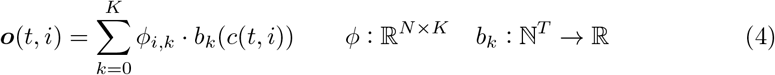

where *K* is the number of learned basis functions. This parameterization allows us to disentangle the forms of individual variability (*individuality basis functions*) and the distribution of this variability (*individuality embeddings*) in the data.

We implement the functions ***m***(*c*(*t, i*)) and ***b***(*c*(*t, i*)) with small fully connected neural networks with non-linearities and several hidden layers. The parameters *θ* of these functions and the entities’ embeddings *ϕ* are learned jointly using minibatch stochastic gradient descent:

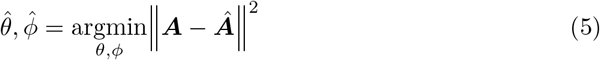

Note that non-negativity is not strictly necessary, but we only consider the non-negative case in this work for consistency with prior work [30, 31]. Furthermore, instead of one common property with discrete time steps, the factors could depend on multiple continuous properties, i.e. *c* : ℝ^*T×N*^, → ℝ^*P*^ e.g. the day and time in a intraday analysis of social networks.

We find that the model’s interpretability can be improved using additional regularization terms without significantly affecting its performance. We encourage sparsity in both the number of used factors and individuality basis functions by adding *L*_1_ penalties of the mean absolute magnitude of the factors ***f***(*t, i*) and basis functions ***b***(*c*(*t, i*)) to the objective. We encourage individuals’ lifetimes to be represented with a sparse embedding using an *L*_1_ penalty of the learned *individuality embeddings ϕ*.

We also introduce an optional adversarial loss term to encourage the model to learn embeddings that are semantically consistent over time, i.e. to only represent two entities that were present in the dataset at different times with different embeddings if this is strictly necessary to factorize the matrices **A**. We jointly train a discriminative network *d*(*ϕ*_*i*_) that tries to classify the time of the first occurrence of all entities based on their *individuality embeddings ϕ*. The negative cross-entropy loss of this model is added as a regularization term to equation 5 in a training regime similar to generative adversarial networks [44]. Note that a high cross-entropy loss of the discriminative network *d*(*ϕ*_*i*_) implies that the distribution of *individuality embeddings ϕ* is consistent over time. See appendix S1.1 for more details and S2 for an ablation study of the effect the individual regularization terms have on the results of the model.

We implemented the model using PyTorch [45] and trained it in minibatches of 256 individuals for 200 000 iterations with the Adam optimizer [46]. We calculate the reconstruction loss ‖**A**_*t*_ − **Â**_*t*_‖^2^ only for valid entries, i.e., we mask out all matrix elements where one of the individuals is not alive at the given time *t*. See appendix S1.3 for the architecture of the learned functions, a precise description of the regularization losses, and further hyperparameters. The code of our reference implementation is publicly available: https://github.com/nebw/temporal_nmf.

### 2.2 Data

#### 2.2.1 Synthetic data

We created synthetic datasets using a generative model of interactions based on a common latent trajectory of factors and groups with structured variations from this trajectory. We compute the number of interactions between two individuals as the dot product of their latent factors and additive Gaussian noise. Using these datasets we can evaluate whether the model successfully converges and is able to correctly identify which individual belongs to which latent group, even in the presence on high amounts of observational noise. While we believe that such a latent structure exists in most complex social systems, it is not directly observable, and thus, for data from a real system, we can only evaluate the model on proxy measures (see section 2.3) that are observable.

We model a common lifetime trajectory of factors using a smoothed Gaussian random walk in ℝ^+^ with *σ*_walk_ = 1 for the steps of the random walk and *σ*_smoothing_ = 10 for the Gaussian smoothing kernel. See Figure 3 a) for one example of a generated lifetime trajectory with three factors. We then randomly create latent groups by creating smoothed Gaussian random walks that define how these groups differ from the common lifetime trajectory. See Figure 3 b) for the lifetime trajectory of one latent group. For each group, we also define different expected mean lifetimes. We set the average lifetime of an entity to 30 days with a standard deviation of 10 days. We then randomly assign 1024 individuals to those latent groups and also assign random dates of emergence and disappearance of these individuals in the dataset. We then compute the individual factor trajectories for each individual, as can be seen in Figure 3 c). Finally, for 100 days of simulated data, we generate interaction matrices by computing the dot products of the factors of all individuals (Figure 3 d).

**Figure 3.**
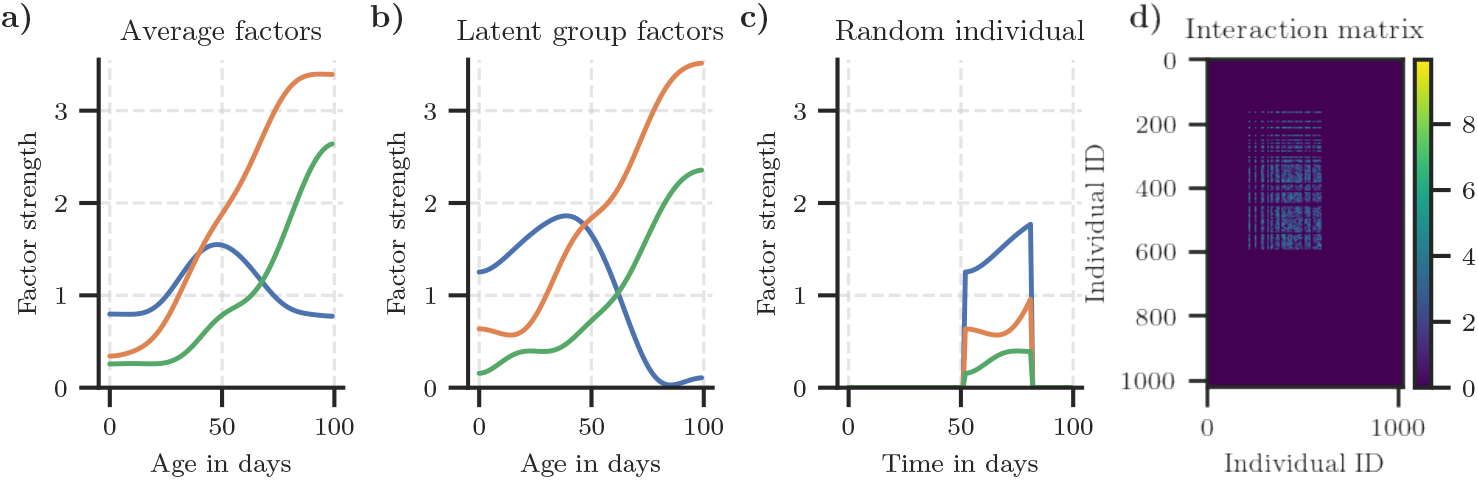
Example of one synthetic dataset. **a)** Common lifetime trajectory of all entities. **b)** The lifetime trajectory of one latent group. **c)** The factors of one individual in the dataset of the latent group visualized in **b. d)** Generated interaction matrix for one day.

We then measure how well the *individuality embeddings ϕ* of a fitted model match the true latent groups from the generative model using the adjusted mutual information score [47]. Furthermore, we measure the mean squared error between the ground truth factors and the best permutation of the factors ***f***^+^. We evaluate the model on 128 different random synthetic datasets with increasing Gaussian noise levels in the interaction tensor.

#### 2.2.2 Honey bee data

Honey bees are an ideal model system with a complex and highly dynamic social structure. The entire colony is observable most of the time. In recent years, technological advances have made it possible to automatically track individuals in entire colonies of honey bees over long periods of time [6,10,48]. We analyze a dataset obtained by tracking thousands of individually marked honey bees at high temporal and spatial resolution, covering entire lifespans and multiple generations.

Two colonies of honey bees were continuously recorded over a total of 155 days. Each individual was manually tagged at emergence, so the date of birth is known for each bee. Timestamps, positions, and unique identifiers of all (N=9286) individuals from these colonies were obtained using the BeesBook tracking system [10, 12, 48]. See Table 1 for dates and number of individuals. Temporal affinity matrices were derived from this data as follows: For each day, counts of proximity contact events were extracted. Two individuals were defined to be in proximity if their markers’ positions had an euclidean distance of less than 2 cm for at least 0.9 seconds. The daily affinity between two individuals *i* and *j* based on their counts of proximity events *p*_*t,i,j*_ at day *t* was then computed as: 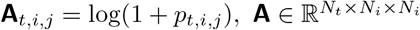, where *N*_*t*_ is the number of days and *N*_*i*_ the number of individuals in the dataset.

**Table 1.** The honey bee datasets contain the number of proximity-inferred interactions extracted from tracking data of all individuals in two long-term recordings spanning a total of 155 days and 9286 individuals.

| Dataset | Dates | Days | Individuals | Interaction pairs |
| --- | --- | --- | --- | --- |
| BN16 | 2016-07-23 to 2016-09-17 | 56 | 2443 | 43 174 748 |
| BN19 | 2019-07-25 to 2019-11-01 | 99 | 6843 | 167 366 381 |

The datasets also contains labels that can be used in proxy tasks (see section 2.3) to quantify if the learned embeddings and factors are semantically meaningful and temporally consistent.

The datasets are open access and available under the *Creative Commons Attribution 4*.*0 International* license: https://zenodo.org/record/3862966 [49].

In both datasets, we define *c*(*t, i*) as the age in days of an individual *i* at time *t*.

### 2.3 Evaluation

#### Reconstruction

We measure how well the original interaction matrices **A** can be reconstructed from the factors. We do not require the model to reconstruct the interaction matrices as well as possible because we only use the reconstruction as a proxy objective to learn a meaningful representation. Still, a high reconstruction loss could indicate problems with the model, such as excessive regularization.

#### Consistency

We measure to what extent the *individuality embeddings ϕ* change over time. For each model, we train a multinomial logistic regression model to predict the source cohort (date of birth) and calculate the area under the ROC curve (AUC_cohort_) using a stratified 100-fold cross-validation with scikit-learn [50]. The baseline models do not learn an individuality embedding; therefore we compute how well the model can predict the cohort using the mean factor representation of the individuals over their lives. We define consistency as 1 − AUC_cohort_ of this linear model. Note that a very low temporal consistency would indicate that the development of individual bees changes strongly between cohorts and colonies, which we know not to be true.

#### Mortality and Rhythmicity

We evaluate how well a linear regression model can predict the mortality (number of days until death) and circadian rhythmicity of the movement [12] (*R*^2^ score of a sine with a period of 24 h fitted to the velocity over a three-day window). These metrics are strongly correlated with an individual’s behavior (e.g. foragers exhibit strong circadian rhythms because they can only forage during the daytime; foragers also have a high mortality). We follow the procedure given in [12] and report the 100-fold cross-validated *R*^2^ scores for these regression tasks.

#### Time spent on different nest substrates

For a subset of the data, from 2016-08-01 to 2016-08-25, nest substrate usage information is also available. This data contains the proportion of time each individual spends in the brood area, honey storage, and on the dance floor. This data was previously published and analyzed [12, 51]. The task of an honey bee worker is strongly associated with her spatial distribution in the hive. We therefore expect a good representation of the individuals’ functional role to correlate with this distribution.

For this data, we expect the factors ***f***^+^ and *individuality embeddings ϕ* to be semantically meaningful and temporally consistent if they reflect an individual’s behavioral metrics (mortality and rhythmicity) and if they do not change strongly over time (measured in the consistency metric).

### 2.4 Baseline models

#### Biological Age

Task allocation in honey bee is partially determined by temporal polyethism. Certain tasks are usually carried out by individuals of about the same age, e.g. young bees are usually occupied with nursing tasks. We therefore use the age of an individual as a baseline descriptor.

#### Symmetric NMF

We compute the factors that optimally reconstruct the original interaction matrices using the standard symmetric NMF algorithm [31, 52], for each day separately, using the same number of factors as in the TNMF model.

#### Optimal permutation SymNMF

We consider a simple extension of the standard SymNMF algorithm that aligns the factors to be more consistent over time. For each pair of subsequent days, we consider all combinatorial reorderings of the factors computed for the second day. For each reordering, we compute the mean *L*_2_ distance of all individuals that were alive on both days. We then select the reordering that minimizes those pairwise *L*_2_ distances and greedily continue with the next pair of days until all factors are aligned. Furthermore, we align the factors across colonies (where individuals cannot overlap) as follows: we run this algorithm for both datasets separately and align the resulting factors by first computing the mean embedding for all individuals grouped by their ages. As before, we now select from all combinatorial possibilities the reordering that minimizes the *L*_2_ distance between the embeddings obtained from both datasets. See section S3.1 for pseudo code.

#### Tensor decomposition

We also compare against a constrained non-negative tensor decomposition model with symmetric factors 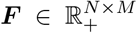 and temporal dynamics constrained to the diagonals, i.e. 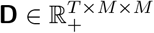 and 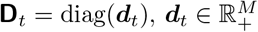.

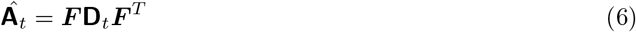

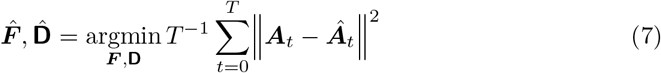

#### Temporal NMF models

We evaluate variants of the temporal symmetric matrix factorization algorithms proposed by [35] and [36].

For the tensor decomposition and temporal NMF baselines, we follow the procedure given above for the *Optimal permutation SymNMF* to find the optimal reordering to align the factors obtained by applying models to the two datasets separately.

## 3 Results

### 3.1 Synthetic data

We factorize the interaction matrices of the 128 synthetic datasets with varying levels of Gaussian noise. We confirmed that our model converges in all datasets and evaluate whether we can distinguish the individuals’ ground truth group assignments. To that end, we extract the *individuality embeddings ϕ* from the models and measure how well they correspond to ground truth data using the adjusted mutual information (AMI) score. Furthermore, we measure the mean squared error between the best permutation of learned factors ***f***^+^ and the ground truth factors.

We find that for low levels of noise, our model can identify the truth group assignments with high accuracy, and are still significantly better than random assignments even at very high levels of noise (see figure 4). Note that for this experiment, we evaluated a model with the same hyperparameters as used in all plots in the results section (see Table 2) and a variant without explicit regularization except the *L*_1_ penalty of the learned *individuality embeddings ϕ* (*λ*_embeddings_, because this regularization is required to meaningfully extract clusters), which was set to 0.1. See appendix 2.2.1 for more details on the synthetic datasets.

**Table 2.**
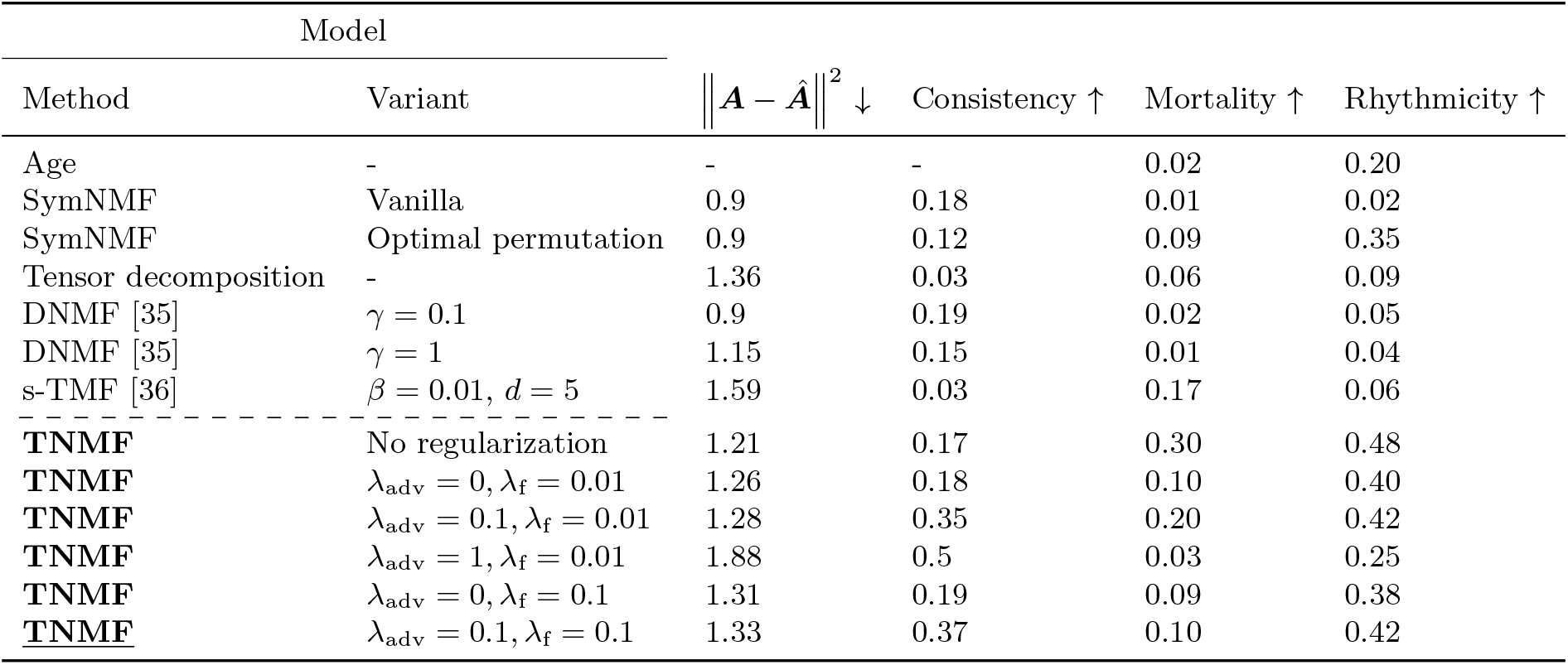
The evaluation metrics for TNMF and the baseline models described in section 2.3. See appendix S1.3 and S3 for descriptions of the hyperparameters used. Note that the SymNMF model reconstruction loss can be seen as a lower bound for the matrix factorization models considered here, and imposing a temporal structure or regularization causes all models to explain less variance in the data. However, for all models except TNMF this does not result in a significant increase of the other metrics. The underlined model is used in all plots in the results section.

| Model |  |  |  |  |  |
| --- | --- | --- | --- | --- | --- |
| Method | Variant | $\ \mathbf{A} - \hat{\mathbf{A}}\ ^2 \downarrow$ | Consistency ↑ | Mortality ↑ | Rhythmicity ↑ |
| Age | - | - | - | 0.02 | 0.20 |
| SymNMF | Vanilla | 0.9 | 0.18 | 0.01 | 0.02 |
| SymNMF | Optimal permutation | 0.9 | 0.12 | 0.09 | 0.35 |
| Tensor decomposition | - | 1.36 | 0.03 | 0.06 | 0.09 |
| DNMF [35] | $\gamma = 0.1$ | 0.9 | 0.19 | 0.02 | 0.05 |
| DNMF [35] | $\gamma = 1$ | 1.15 | 0.15 | 0.01 | 0.04 |
| s-TMF [36] | $\beta = 0.01, d = 5$ | 1.59 | 0.03 | 0.17 | 0.06 |
| **TNMF** | No regularization | 1.21 | 0.17 | 0.30 | 0.48 |
| **TNMF** | $\lambda_{\text{adv}} = 0, \lambda_{\text{f}} = 0.01$ | 1.26 | 0.18 | 0.10 | 0.40 |
| **TNMF** | $\lambda_{\text{adv}} = 0.1, \lambda_{\text{f}} = 0.01$ | 1.28 | 0.35 | 0.20 | 0.42 |
| **TNMF** | $\lambda_{\text{adv}} = 1, \lambda_{\text{f}} = 0.01$ | 1.88 | 0.5 | 0.03 | 0.25 |
| **TNMF** | $\lambda_{\text{adv}} = 0, \lambda_{\text{f}} = 0.1$ | 1.31 | 0.19 | 0.09 | 0.38 |
| **<u>TNMF</u>** | $\lambda_{\text{adv}} = 0.1, \lambda_{\text{f}} = 0.1$ | 1.33 | 0.37 | 0.10 | 0.42 |

**Figure 4.**
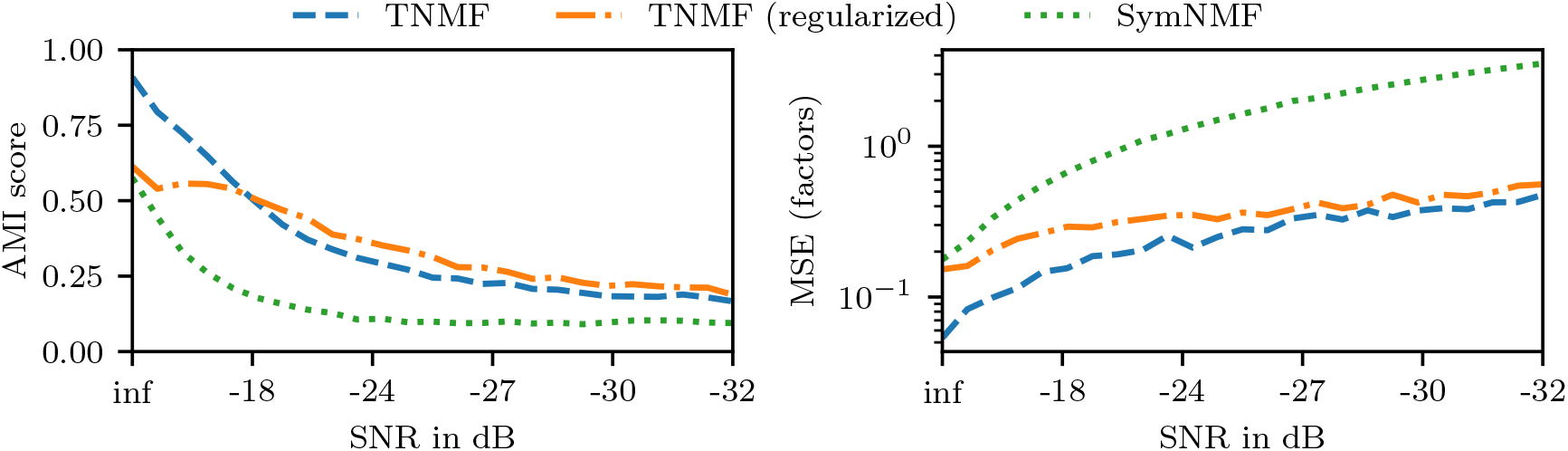
AMI score and mean squared error between true factors and the best permutation of learned factors for increasing noise levels. The median values over 128 trial runs are shown.

### 3.2 Honey bees

#### Mean lifetime model

The model learns a sparse representation of the developmental trajectory of a honey bee in the space of social interactions. Only two factors are effectively used (they exceed the threshold value of 0.01). These factors show a clear trend over the life of a bee, indicating that the model captures the temporal aspects of the honey bee division of labor (See Figure 5).

**Figure 5.**
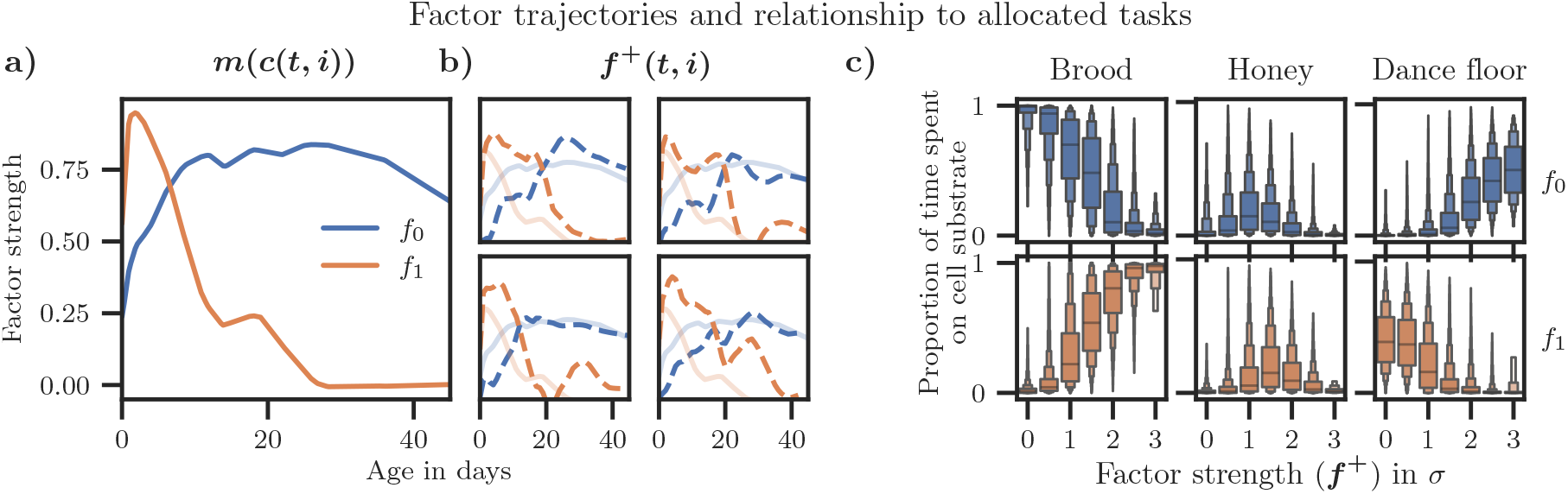
**a)** Mean lifetime trajectories according to ***m***(*c*(*t, i*)). The model learns a sparse representation of the functional position of the individuals in the social network. *f*_0_ (blue) mostly corresponds to middleaged and older bees, and *f*_1_ (orange) predominantly describes young bees. Only factors with a mean magnitude of at least 0.01 are shown. **b)** Even though the model uses only these two factors, it is still expressive enough to capture individual variability, as can be seen in randomly sampled individuals’ lifetime trajectories. **c)** The individual factors ***f***^+^ and the proportion of time the individuals spent on different nest substrates. The strong correlation indicates that the learned factors are a good representation of the individuals’ roles in the colonies. Note that the factors have been divided by their standard deviation here for ease of comparability.

#### Interpretability of factors

To understand the relationship between the factors and division of labor, we calculate how the factors map to the fraction of time an individual spent on the brood area, honey storage, or dance floor (where foragers aggregate). Time spent on these different substrates is a strong indicator of an individual’s task. The factor *f*_1_, which peaks at young age (Figure 5), correlates with the proportion of time spent in the brood area, while a high *f*_0_ indicates increased time spent on the dance floor. Therefore, the model learned to map biologically relevant processes.

#### Individuality basis functions and individuality embeddings

Due to the regularization of the embeddings, the model learns a sparse set of *individuality basis functions*. As encouraged by the model, most individuals can predominantly be described by a single basis function. That means that while each honey bee can collect a unique set of experiences, most can be described with a few common *individuality embeddings* which are consistent across cohorts and colonies. In the context of honey bee division of labor, the basis functions are interpretable because the factors correspond to different task groups. For example, *b*_12_(*c*(*t, i*)) (accounting for ≈10.7% of the individuals) describes workers that occupy nursing tasks much longer than most bees. As the *individuality embeddings ϕ* only scale the magnitude of the basis functions, they can be interpreted in the same way. *Individual lifetime trajectories* in the factor space can be computed based on the *mean lifetime trajectories* (***m***), *individuality basis functions* (***b***(*c*(*t, i*))) and *individuality embeddings* (*ϕ*). See figure 6 for examples of *individual lifetime trajectories* from workers that most strongly corresponded to the common *individuality basis functions*.

**Figure 6.**
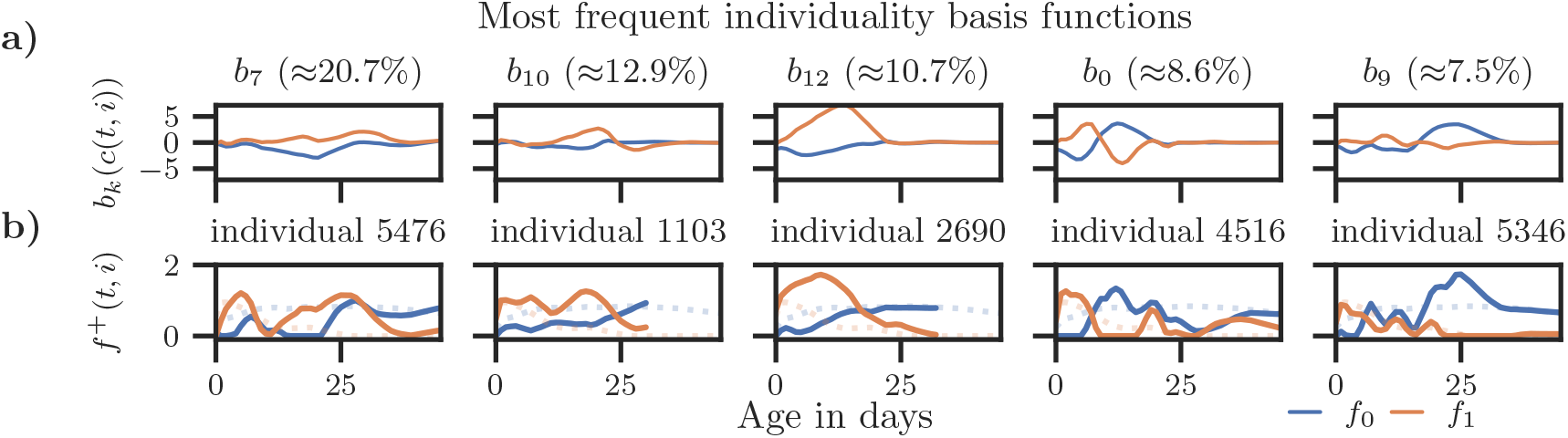
**a)** Magnitude of factor offsets for the five most common individuality basis functions over age *b*_*k*_(*c*(*t, i*)). The percentage of individuals that most strongly correspond to the individual basis functions is shown in the column titles. More than 60% of the individuals strongly correspond to one of the five basis functions shown here. **b)** Because the basis functions describe *individuality offsets* from the mean lifetime trajectory, it may be easier to interpret them by visualizing individual examples. For each of the basis functions (top row), we show a lifetime trajectory of an individual that corresponds to that basis function (bottom row). Note that individuals can die or disappear at any time (solid lines). The mean lifetime trajectories are shown as dotted lines in the background.

#### Evaluation

We verify that the learned representations of the individuals are meaningful (i.e., they relate to other properties of the individuals, not just their interaction matrices) and semantically consistent over time and across datasets using the metrics described in the section *Evaluation*. We compare variants of our model with different adversarial loss scaling factors and factor *L*_1_ regularizations, the baseline models, and the individuals’ ages. We expect a good model to be temporally consistent and semantically meaningful. All variants of our model outperform the baselines in terms of the semantic metrics *Mortality* and *Rhythmicity*, except for the [36] model, which performs comparably well in the *Mortality* metric. The adversarial loss term further increases the *Consistency* metric without negatively affecting the other metrics. A very strong adversarial regularization (see row with *λ*_adv_ = 1 in Table 2) prevents the model from learning a good representation of the data. See Table 2 for an overview of the results. We also evaluate the tradeoff between the different metrics using a grid search over the hyperparameters (see appendix 3.2).

#### Scalability

The functions ***m***(*c*(*t, i*)) and ***b***(*c*(*t, i*)) are learned neural networks with non-linearities. The objective is non-convex and we learn the model parameters using stochastic gradient descent. Optimization is therefore slower than the standard NMF algorithms that can be fitted using algorithms such as Alternating Least Squares [53]. We found that the model converges faster if the reconstruction loss of the age based model ***m***(*c*(*t, i*)) is additionally minimized with the main objective in equation 5. Due to the minibatch training regime, our method should scale well in larger datasets. Small neural networks were sufficient to learn the functions ***m***(*c*(*t, i*)) and ***b***(*c*(*t, i*)) in our experiments. Most of the runtime during training is spent on the matrix multiplication *f*^+^(*t, i*) · *f*^+^(*t, j*)^*T*^ and the corresponding backwards pass.

#### Tradeoff between temporal consistency and semantic meaningfulness

We performed a grid search over the hyperparameters *λ*_f_, *λ*_adv_, *λ*_basis_, and *λ*_embeddings_ (see Table 1) to evaluate whether models can only be either semantically meaningful or temporally consistent. For this analysis, we define *Semantic meaningfulness* as the sum of the *Rhythmicity* and *Mortality* metrics introduced in section 2.3. We find that models that are very temporally consistent fail to learn semantically meaningful information. Interestingly, the models with the best tradeoff between the two metrics are almost as semantically meaningful as those models with low temporal consistency and the highest semantic meaningfulness. This analysis suggests that regularization encourages the model to only represent different individuals differently if this is strictly necessary to factorize the data. See Figure 5.

## 4 Discussion

Temporal NMF factorizes temporal matrices with overlapping and even disjoint communities by learning an embedding of individuals as a function of a common factor, such as age, and a learned representation of the individuals’ individuality. This explicit dependency on a common factor that partially determines the function of an individual constitutes an inductive bias. We show that the model learns semantically consistent representations of individuals, even in challenging cases, such as the datasets analyzed in this work.

The individual components of the model are straightforward to visualize and interpret. The learned individuality embeddings *ϕ* can be understood as soft-cluster assignments relating to the whole lifetime of an individual, while the factor vectors ***f***^+^(*t, i*) can be interpreted as cluster assignments of the individuals at a specific point in time, i.e. two individuals with similar factor vectors are likely to interact if they exist in the same group at the same time. Furthermore, the model encourages sparsity, making the results easier to interpret because the model only uses as many factors and clusters as necessary.

We identified a crucial trade-off that comes with temporal consistency: For a specific point in time, the ability to predict behaviorally relevant attributes will likely be worse for a model that learns temporally consistent representations compared to a non-consistent model with the same capacity. Conversely, in more challenging cases, e.g. when taking long periods of time or data from disjoint communities into consideration, temporally consistency is indispensable for a good representation. Furthermore, we found that models can be temporally consistent, semantically meaningful, or both; selecting the correct model requires an inductive bias, but regularization of the model also influences the results.

Previous works have demonstrated that biologically relevant findings can be obtained using network analysis of social interaction networks [4, 5, 26, 54–62]. A recent method, *Network Age* [12], proposes using spectral decomposition of honey bee interaction networks into succinct descriptors of the individual’s social network that can be used to predict task allocation, survival, activity patterns, and future behavior. Symmetric nonnegative matrix factorization and Laplacian-based spectral clustering have been shown to be equivalent [29]. Thus, TNMF can be understood as a further development of *Network Age*. TNMF learns representations of individuals based on their social interaction network that can facilitate the analysis of developmental trajectories, division of labor, and individual variance in behavior. Furthermore, TNMF provides temporally consistent embeddings and with that rectifies a remaining limitation of *Network Age*. We confirmed that, on the honey bee dataset, TNMF obtains biologically meaningful lifetime trajectories with promising prospects for experimental application. TNMF may help advance our understanding of the colony function and the interplay between environmental factors and individual and collective responses. The method presented here offers a way to investigate the impact of stress factors, such as pesticides, parasitic mites, and agricultural monoculture, on the social structure of colonies and therefore may present an avenue to improve honeybee health.

**Figure 7.**
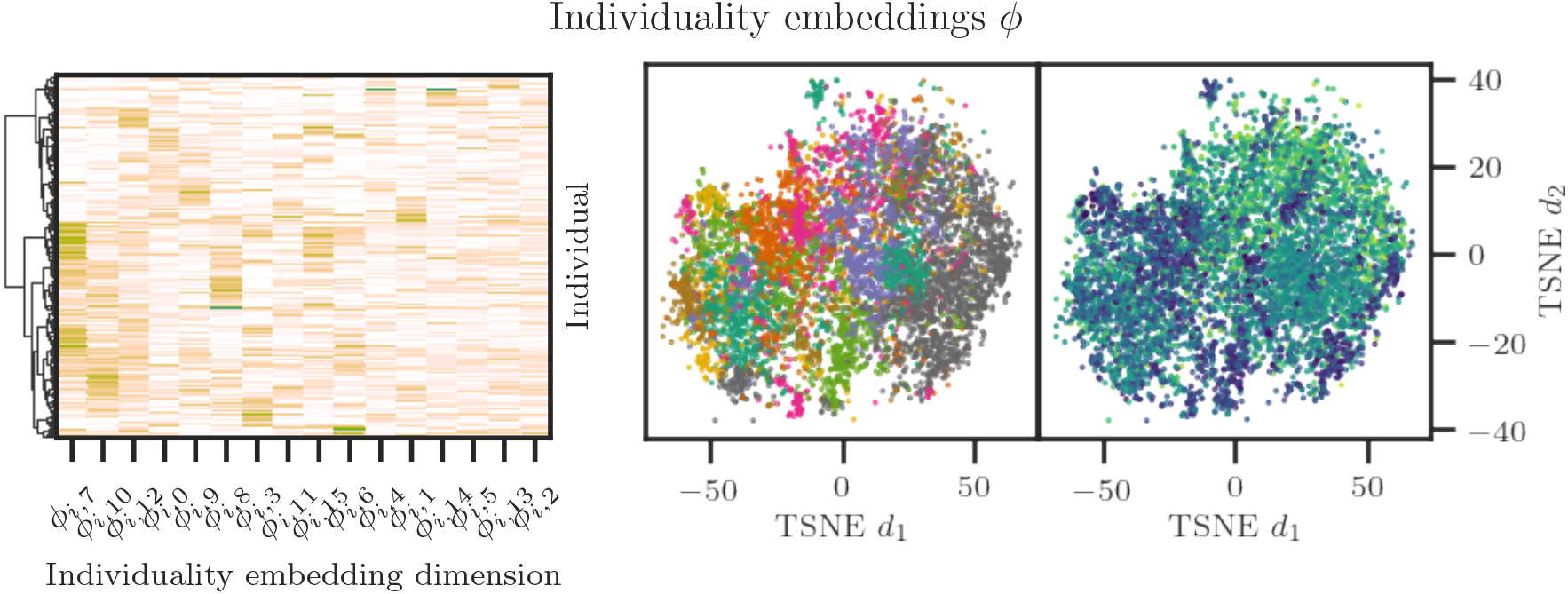
**Left:** Hierarchical clustering of individuality embeddings: Most individuals strongly correspond to a single individuality basis function, making it easy to cluster their lifetime social behavior (i.e. each individual has a high value in a single dimension for their individuality embedding). Because each cluster is strongly associated with a specific individuality basis function, and because each basis function is interpretable (Figure 5), these blueprints of lifetime development can also be intuitively understood and compared. **Right:** TSNE plots of the individuality embeddings colored by cluster (left) and the maximum circadian rhythmicity of an individual during her lifetime (right), indicating that the embeddings are semantically meaningful.

We applied our method to the honey bee model, which has numerous individuals, with an entangled and highly dynamic social structure. Our method, however, can be applied to any setting in which matrix factorization is commonly used, such as recommender systems, network analysis, audio processing, bioinformatics, etc. Interaction matrices in networked systems have a broad class of use-cases. In any system with dynamically interacting units, our model reduces high-dimensional interaction patterns to low-dimensional embeddings. The only requirement is that the interactions follow our generic model of an average path from which individual units can deviate. The method could serve as a means to generate hypotheses: Clustering individuals in the embedding space may reveal functional groups, and the basis functions can indicate relevant time points in individual developments that can be investigated in follow-up studies. Note that time in our model may be replaced by any variable along which one wants to study the matrix dynamics. While we evaluate TNMF on honey bees, the method may be used to study human social networks and their underlying dynamics. A deeper understanding of human interaction dynamics may benefit aspects of human life, such as health, technology, and work. The method could, for example, be used to identify individuals with a higher risk of contracting or transmitting a disease, or help assess the effect of pandemics and potential interventions.

By publishing the honey bee dataset and our reference implementation of the TNMF algorithm, we hope to encourage the scientific community to build upon our efforts.

## Acknowledgments

We thank Leon Sixt for valuable discussions and comments on the manuscript, and numerous students for assisting with data aquisition and preprocessing in the BeesBook system.

## Notes

https://zenodo.org/record/3862966

### Competing Interest Statement

The authors have declared no competing interest.

