## Supporting Information for "Learning to embed lifetime social behavior from interaction dynamics"

### S1 Model details and hyperparameters

#### S1.1 Regularization terms

$$R_{\text{embeddings}} = \lambda_{\text{embeddings}} N_i^{-1} \sum_{i=0}^{N_i} \sum_{k=0}^{K} \left| \phi_{i,k} \right| \tag{8}$$

$$R_{\text{f}} = \lambda_{\text{f}} N_i^{-1} \sum_{i=0}^{N_i} \sum_{t=0}^{N_t} f^{+}(t,i) \tag{9}$$

$$R_{\text{basis}} = \lambda_{\text{basis}} N_a^{-1} \sum_{a=0}^{N_a} \sum_{k=0}^{K} \left| b_k(a) \right| \quad N_a = 60 \tag{10}$$

where $N_a$ can be any number higher than the oldest individual in the dataset at any time.

$$R_{\text{adv}} = \lambda_{\text{adv}} N_i^{-1} \sum_{i=0}^{N_i} \log \left( \frac{\exp(\boldsymbol{d}(\phi_i)_{c[i]})}{\sum_{d}^{N_d} \exp(\boldsymbol{d}(\phi_i)_d)} \right) \tag{11}$$

where $\boldsymbol{d}(\phi_i)$ is the probability distribution returned by the discriminative network, $c[i]$ the day the entitity $i$ emerged in the dataset, and $N_d$ the number of days in the dataset.

See appendix S2 for an ablation study of the effect of these regularization term on the results.

#### S1.2 Network architecture

We use the following neural network architecture for the functions $\boldsymbol{m}(c(t,i))$, $\boldsymbol{b}(c(t,i))$, and $\boldsymbol{d}(\phi_i)$:

$$\text{Linear}(N_{in}, N_h) \to \text{LReLU} \to \underbrace{\text{Linear}(N_h, N_h) \to \text{LReLU}}_{N_l\text{-times}} \to \text{Linear}(N_h, N_{out})$$

where *Linear* is an affine transformation $f(x) = Ax + b$ and $\alpha = 0.3$ for the Leaky ReLU activation function. For $\boldsymbol{m}(c(t,i))$ and $\boldsymbol{b}(c(t,i))$: $N_{in} = 1$ (the individuals' ages). For $\boldsymbol{m}(c(t,i))$: $N_{out} = M$ and for $\boldsymbol{b}(c(t,i))$: $N_{out} = MK$. For $d(\phi_i)$: $N_{in} = K$ and $N_{out} = N_{\text{labels}}$.

#### S1.3 Hyperparameters

The scaling factors for the regularization losses (see Table 1) were manually selected by increasing each factor until it prevented the model from converging (i.e. the reconstruction loss of the full model $\boldsymbol{f}^{+}(t,i)$ did not improve on the age model $\boldsymbol{m}$). This initial set of hyperparameters was then manually refined such that each regularization loss was still effective (e.g. the factor regularization loss $L_f$ reduced the total number of factors effectively used by the model). Overfitting was not a concern because the model is fitted unsupervised and the goal of the hyperparameter selection was to find a set of parameters that is sparse and interpretable, and not to increase the predictive capabilities of the learned factors.

**Supporting Table 1.** Hyperparameters used in the evaluated models (if not stated otherwise)

| Parameter | Value | Description |
|---|---|---|
| $N_l$ | 3 | Number of hidden layers |
| $N_h$ | 64 | Hidden layer size |
| $M$ | 8 | Number of factors |
| $K$ | 16 | Number of individuality basis function |
| $N_{\text{labels}}$ | 100 | Number of cohorts |
| $N_{\text{batch}}$ | 128 | Minibatch size |
| $N_{\text{steps}}$ | 100 000 | Number of training iterations |
| $\lambda_{\text{f}}$ | 0.1 | Factor $L_1$ regularization |
| $\lambda_{\text{adv}}$ | 0.1 | Factor $L_1$ regularization |
| $\lambda_{\text{basis}}$ | 0.01 | Basis function $L_1$ regularization |
| $\lambda_{\text{embeddings}}$ | 0.1 | Embedding $L_1$ regularization |

#### S1.4 Model fitting

The model was fitted using a single GPU (GeForce RTX 2080 Ti). A training run consisting of 200 000 minibatches finished in about six hours. Due to overhead in data loading and preprocessing, up to three training runs could be executed in parallel without negatively affecting the runtime.

**Supporting Algorithm 1:** Training loop

**for** $b = 0$ **to** $N_{steps}$ **do**
  Draw minibatch of $N_{\text{batch}}$ random individuals
  Compute $\hat{\mathbf{A}}_{t,i,j}$ $\forall t$ for individuals $i, j$ in minibatch
  Compute model training loss: equation 5 + regularization
  Update parameters for $\boldsymbol{m}(c(t,i))$, $\boldsymbol{b}(c(t,i))$, and $\boldsymbol{d}(\phi_i)$
  Compute $\hat{l}_i = \boldsymbol{d}(\phi_i)$ for individuals $i$ in minibatch
  Compute discriminator training loss
  Update parameters for $\boldsymbol{d}(\phi_i)$
**end for**

### S2 Ablation study

As highlighted in section 3, the regularization terms $R_{\mathrm{f}}$ and $R_{\mathrm{adv}}$ improve the semantic meaningfulness and temporal consistency of the model. Here we present an ablation study that shows the effects of the regularization terms on the sparseness and interpretability of the learned factors and embeddings, and how the adversarial term influences the distribution of the learned embeddings. While the regularization terms increase the methodological complexity of the model, we argue that they improve the interpretability of the results.

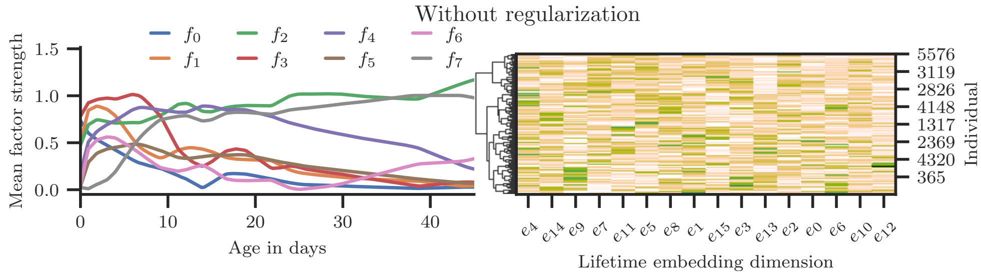


**Supporting Figure 1.** A model trained without the regularization terms $R_{\mathrm{embeddings}}$, $R_{\mathrm{f}}$, $R_{\mathrm{basis}}$, and $R_{\mathrm{adv}}$. The model uses all eight factors, and many of them are strongly correlated (left). Most individuals personality offsets are a function of multiple embeddings (right).

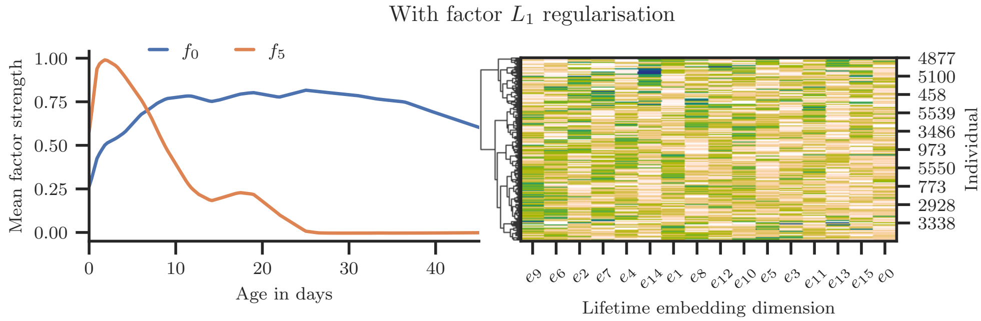


**Supporting Figure 2.** A model trained with only the regularization term $R_{\mathrm{f}}$. The factor trajectories are now sparse and uncorrelated (left), most individuals personality offsets are a still a function of multiple embeddings (right).

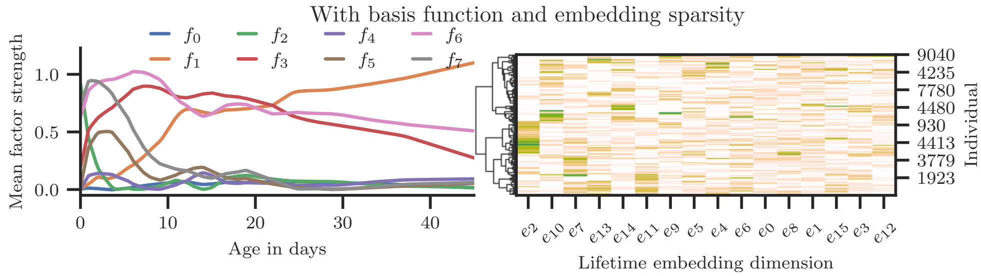


**Supporting Figure 3.** The regularization terms $R_{\mathrm{embeddings}}$ and $R_{\mathrm{basis}}$ introduce sparseness in the embeddings (right), and also slightly decorrelate the factor trajectories (left).

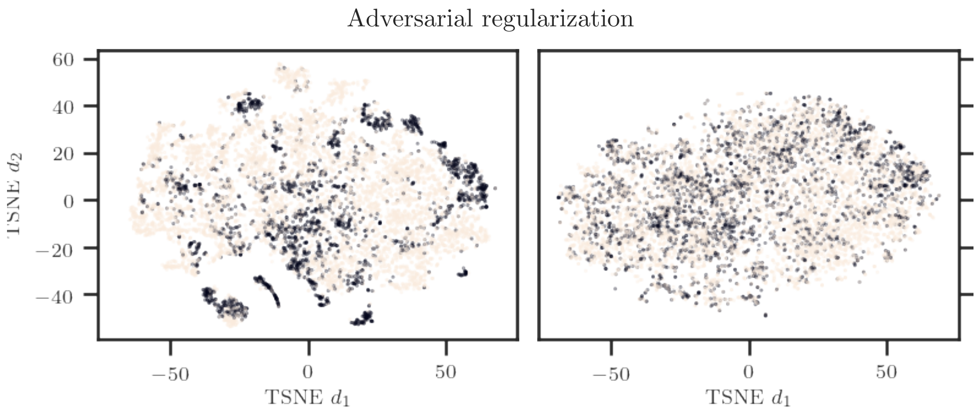


**Supporting Figure 4.** Scatter plots of the individuality embeddings $\phi$ (reduced to two dimensions using TSNE) for two models trained without (left) and with adversarial (right) regularization. The color encodes the dataset of the individuals (Dark = BN16, Bright = BN19). The model with adversarial regularization learns to embed the individuals from two different colonies that never interacted with each other in a joint individuality embedding space.

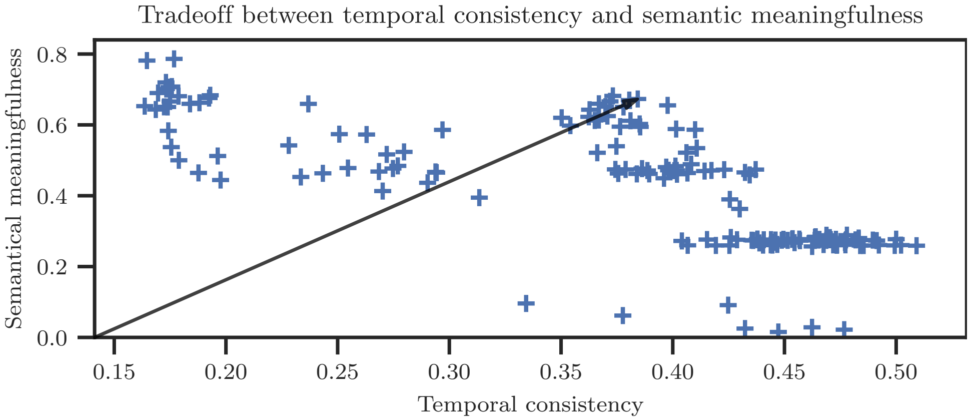


**Supporting Figure 5.** A grid search over the hyperparameters reveals that models can be temporally consistent, semantically meaningful, or both. Semantic meaningfulness is the sum of the *Rhythmicity* and *Mortality* metrics introduced in section 2.3. The arrow points to the model with the best tradeoff.

### S3 Baseline models

We implemented SymNMF, and models proposed in [35] and [36] in PyTorch and compare them to our method. Following the notation given in [36], we list the degree of the fitted polynomial as $d$, and the regularization parameter as $\beta$. For the model proposed by [35], we list the regularization as $\gamma$. For SymNMF, the *Aligned* variant refers to the *Aligned symmetric NMF* described in section 2.4.

We only evaluate the non-negative and symmetric variants of the models for consistency with our method. For all baselines, we only list the hyperparameters with the best results that we were able to obtain.

#### S3.1 Temporal reordering of factors for baseline models

**Supporting Algorithm 2:** Temporal reordering of factors for baseline models

**for** $d = 0$ **to** $N_{days}$ **do**
  $F_{\text{result}} < -$
  **if** $d == 0$ **then**
    $F_{\text{previous\_reordered}} \leftarrow$ Precomputed SymNMF factors of day 0
  **else**
    $F_{\text{current}} \leftarrow$ Precomputed SymNMF factors of day $d$
    $\text{min\_loss} \leftarrow 0$
    $F_{\text{best}} \leftarrow F_{\text{current}}$
    **for** all permutations $p$ of orderings of factors $[0..M]$ **do**
      $F_{\text{current\_reordered}} \leftarrow F_{\text{current}}$ reordered by permutation $p$
      **if** $(F_{\text{current\_reordered}} - F_{\text{previous\_reordered}})^2 < \text{min\_loss}$ **then**
        $\text{min\_loss} \leftarrow (F_{\text{current\_reordered}} - F_{\text{previous\_reordered}})^2$
        $F_{\text{best}} \leftarrow F_{\text{current\_reordered}}$
      **end if**
    **end for**
    $F_{\text{previous\_reordered}} \leftarrow F_{\text{best}}$
  **end if**
  $F_{\text{result}}.insert(F_{\text{previous\_reordered}})$
**end for**
**return** $F_{\text{result}}$

### S4 Learned basis functions and individual trajectories

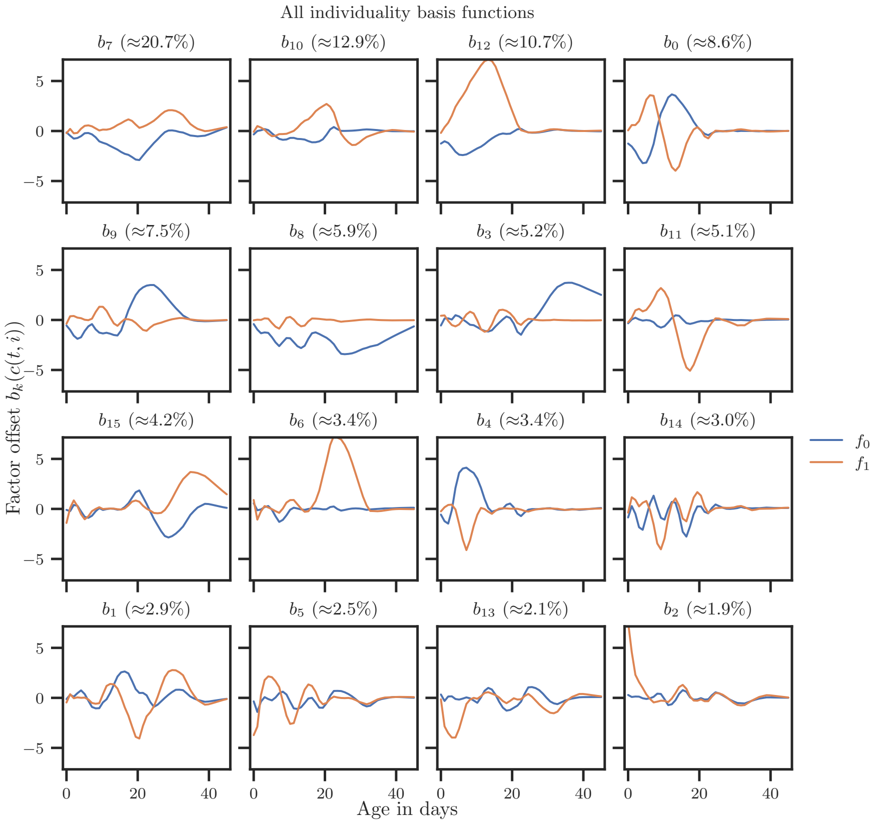


**Supporting Figure 6.** Magnitude of factor offsets for all learned individuality basis functions over age $b_k(c(t,i))$.

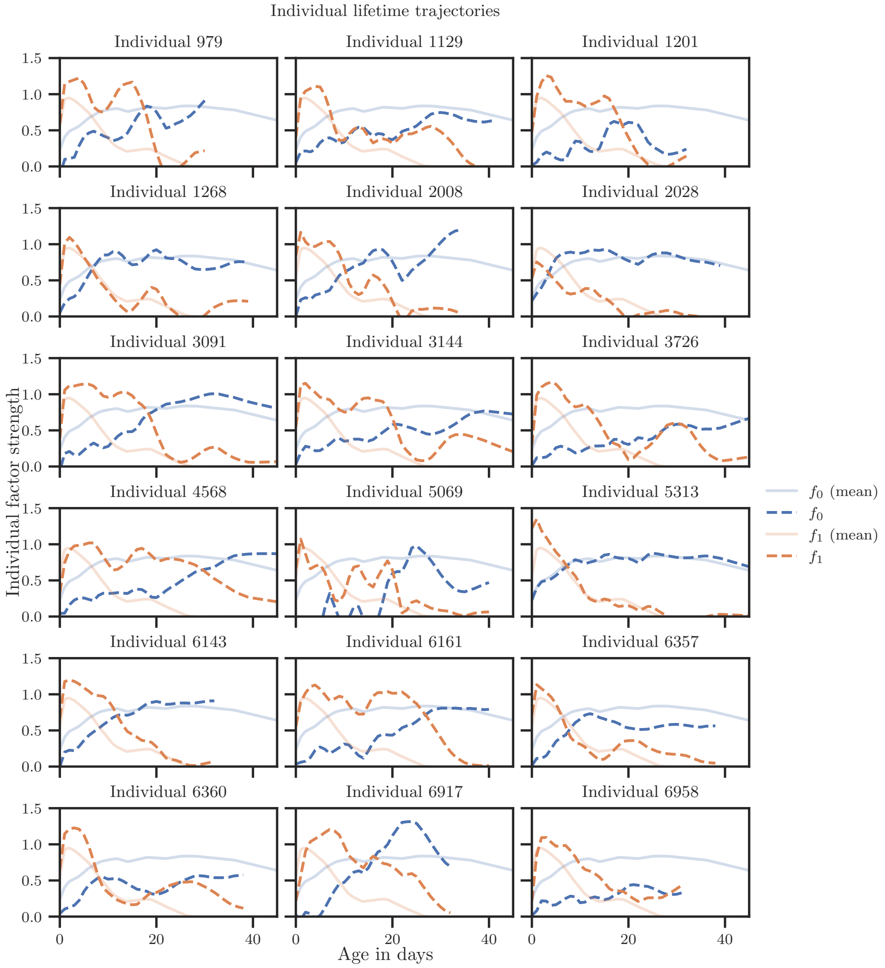


**Supporting Figure 7.** Individual lifetime trajectories: 18 individuals that lived for at least 35 days were randomly sampled and their factors $f(t, i)$ were computed over all $t$, constituting their lifetime trajectories.
